# Mesoscale Heterogeneity Shapes Molecular Selectivity and Transport in Biomolecular Condensates

**DOI:** 10.64898/2026.09.05.749622

**Authors:** Vladimir Grigorev, Yaojun Zhang

**Author notes:** (YZ). Contact authors (VG).

## Abstract

Growing experimental and computational evidence suggests that many condensates exhibit heterogeneous internal organization at mesoscopic length scales. How such mesoscale organization influences client selectivity and transport, however, remains poorly understood. Here, we use coarsegrained molecular dynamics simulations to compare homogeneous and heterogeneous sticker-spacer condensates with matched scaffold density, sticker fraction, and sticker-sticker interaction strength. We find that the two condensates exhibit markedly different client partitioning profiles and transport dynamics. In the heterogeneous condensate, clients sample distinct local microenvironments in a size-dependent manner, leading to pronounced differences in their transfer free energy compared with the homogeneous condensate. The heterogeneous internal organization also generates local environments with different mobilities and constraints, giving rise to transient subdiffusion and non-Gaussian displacement distributions in client motion. Our results identify mesoscale organization as an important physical factor controlling both molecular selectivity and transport in biomolecular condensates, with implications for the biochemical functions of both natural and engineered systems.

## I. INTRODUCTION

Biomolecular condensates are dynamic, membrane-less compartments that often form through liquid-liquid phase separation driven by multivalent interactions among proteins and nucleic acids [1–3]. These compartments organize cellular biochemistry via the selective recruitment of molecules from their surroundings [4, 5]. By controlling molecular partitioning, condensates can create biochemical environments that regulate transcription [6–8], signaling [9, 10], stress responses [11–13], and other essential cellular processes. Consequently, dysregulation of condensate composition or dynamics has been implicated in diseases such as neurodegeneration and cancer [14–17]. How condensates achieve their compositional selectivity, however, remains incompletely understood.

Condensates in living cells can contain hundreds of distinct molecular species [18, 19]. To organize their components, condensates generally employ a scaffold-client framework [18, 20]. Scaffolds are the molecules that drive phase separation, whereas clients are recruited into condensates without being necessary for condensate formation. Although typically present at lower concentrations than scaffolds, clients are an integral part of condensates and play key roles in condensate functions [5, 18]. Understanding how clients are selectively incorporated and how their transport is regulated within condensates is therefore essential for understanding condensate function.

Client recruitment is commonly attributed to favorable interactions with scaffold molecules [18, 20, 21], but for successful recruitment, clients must also overcome the physical cost of insertion into a dense environment [22– 25]. Recent work has shown that the conformational entropy of intrinsically disordered scaffold regions can impose a size-dependent exclusion barrier, such that large particles are depleted from condensates unless this barrier is compensated by favorable binding interactions [22].

The balance between entropic exclusion and favorable binding provides a useful picture for understanding client recruitment. However, previous studies following this picture have generally considered spatially uniform dense phases. In many condensates, molecules are not homogeneously mixed but instead form spatially heterogeneous structures at mesoscopic length scales [26–34]. For instance, single-fluorogen imaging has revealed non-uniform localization of environmentally sensitive probes in disordered protein condensates, suggesting the presence of hydrophobic nanoscale hubs that potentially arise from interactions among locally clustered aromatic sticker residues [29]. Complementing these observations, simulations of intrinsically disordered proteins (IDPs) have shown that sequence patterning can drive microphase separation, leading to sticker-rich and spacer-rich domains within the dense phase [28, 32]. Spatial heterogeneity has further been proposed to underlie dynamically heterogeneous diffusion in fused in sarcoma (FUS) condensates [33] and in condensates that form at postsynaptic densities [34]. Together, these observations raise the question of how spatial heterogeneity shapes client partitioning and transport in biomolecular condensates.

Here, we address this question using coarse-grained molecular dynamics simulations and theoretical analysis. We compare a homogeneous condensate formed by alternating sticker-spacer polymers with a heterogeneous condensate formed by blocky polymers that microphase-separate into sticker-rich and spacer-rich domains. The two condensates exhibit markedly different size-dependent partitioning profiles for clients at various affinities to stickers. We show that this difference arises from the distinct local microenvironments within the heterogeneous condensate, which redistribute clients between sticker-rich and spacer-rich regions in a size-dependent manner. These microenvironments also give rise to distinct client mobility states, resulting in transient subdiffusion and non-Gaussian displacement distributions. Thus, mesoscale spatial organization shapes both molecular selectivity and transport in biomolecular condensates.

## II. RESULTS

### A. Spatial heterogeneity shapes molecular selectivity in model condensates

To construct a minimal model of condensates with distinct internal spatial organization, we simulated stickerspacer polymers composed of attractive sticker beads and repulsive spacer beads connected by harmonic springs [35, 36]. We considered two sequences with identical length and sticker fraction: an alternating sequence, (A_1_B_1_)_64_, and a blocky sequence, (A_16_B_16_)_4_, where A and B denote spacer and sticker beads, respectively. The alternating sequence forms a spatially homogeneous dense phase (Fig. 1a), whereas the blocky sequence forms a heterogeneous dense phase with mesoscale organization characterized by sticker-rich and spacer-rich domains [28] (Fig. 1b).

**FIG. 1.**
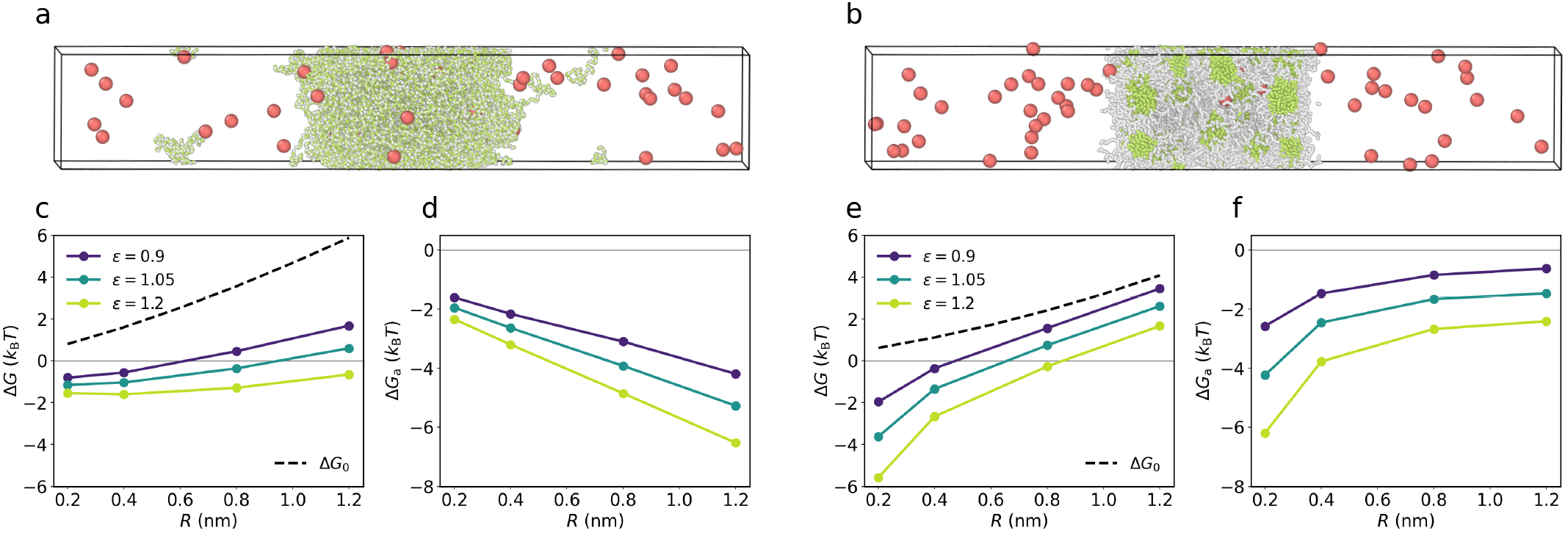
Mesoscale heterogeneity shapes client partitioning in model condensates. **a**, Snapshot of a coarse-grained moleculardynamics simulation of 200 (A_1_B_1_)_64_ polymers together with 50 spherical client particles of type C (red). A and B denote spacer (white) and sticker (green) beads, respectively, both with diameter *σ* = 0.6 nm and are connected by harmonic bonds with equilibrium length *r*_b_ = 0.38 nm. The client particles have radius *R* = 1.2 nm and affinity to stickers *ϵ*_BC_ = 1.2 *k*_B_*T* . The polymers macrophase separate in a 120 nm *×* 20 nm *×* 20 nm simulation box in a slab geometry with periodic boundary conditions, yielding a homogeneous condensate with clients partitioned between the dense and dilute phases (*P* = 1.93). **b**, Same as **a**, but for 200 (A_16_B_16_)_4_ blocky polymers. The polymers microphase separate into sticker-rich and spacer-rich domains, resulting in a spatially heterogeneous condensate, in which the same clients partition to a much lesser extent (*P* = 0.19). **c** and **e**, Total transfer free energy Δ*G* (solid lines) and repulsive contribution Δ*G*_0_ (dashed lines) as functions of client radius *R* at different client-sticker affinities *ϵ*_BC_ for the homogeneous (**c**) and heterogeneous (**e**) condensates. **d** and **f**, Corresponding attractive contribution Δ*G*_a_ = Δ*G* − Δ*G*_0_ for the homogeneous (**d**) and heterogeneous (**f** ) condensates. See Methods for details of the simulations and analysis.

Previous work has shown that client partitioning depends on the dense-phase volume fraction [22]. To disentangle the effect of internal spatial organization from that of overall density, we first determined the phase diagrams for both sequences and selected a sticker-sticker interaction strength of *ϵ*_BB_ = 0.84 *k*_B_*T*, at which the two condensates have the same dense-phase volume fraction *ϕ* = 0.19 (Fig. S1). We then introduced spherical particles of type C as a minimal model of globular clients (Fig. 1a,b). Each particle has a radius *R* and an affinity to stickers *ϵ*_BC_, allowing us to investigate how client partitioning into the two condensates depends on both its size and affinity to stickers. See Methods for simulation details.

We quantified client partitioning using the partition coefficient *P* ≡ *c*_den_*/c*_dil_, where *c*_den_ and *c*_dil_ are the particle concentrations in the dense and dilute phases, respectively. The corresponding transfer free energy from the dilute phase to the dense phase is

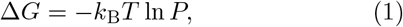

where a positive value of Δ*G* indicates exclusion, whereas a negative value indicates recruitment (Fig. 1c, e). We further decomposed the transfer free energy Δ*G* into a purely repulsive contribution, Δ*G*_0_, arising from excluded volume and the conformational entropy of polymer chains [22] (Fig. 1c, e, dashed lines), and an attractive contribution, Δ*G*_a_, arising from client-sticker attractions (Fig. 1d, f). See Methods for details of the freeenergy quantification.

Despite having the same overall scaffold density and sticker fraction, the two condensates exhibit markedly different size-dependent partitioning profiles (Fig. 1c–f). In the heterogeneous condensate, small adhesive clients are recruited more strongly, whereas larger clients become increasingly excluded compared with the homogeneous condensate (Fig. 1c,e). This enhanced size selectivity reflects differences in the repulsive and attractive contributions to the transfer free energy between the two condensates. The repulsive contribution increases with client size in both condensates. For a given client size, however, |Δ*G*_0_| is lower in the heterogeneous condensate (Fig. 1c,e), reflecting the presence of low-density spacer-rich regions that reduce the cost of particle insertion. Strikingly, the attractive contribution shows opposite size dependence in the two condensates. In the homogeneous condensate, |Δ*G*_a_| grows approximately linearly with client size (Fig. 1d), consistent with previous studies of attractive particle partitioning into homopolymer melts [37]. In contrast, Δ*G*_a_ decreases nonlinearly with client size in the heterogeneous condensate (Fig. 1f). This raises the question of why larger adhesive particles experience weaker attraction in the heterogeneous condensate despite having a larger surface area available for interactions.

To address this question, we characterized the local environment for each client by counting the number of nearby stickers, *n*_st_, and spacers, *n*_sp_, within a distance *r*_c_ = *R*+ 2.5*σ* from the client center, and constructed the joint probability distributions, *p*(*n*_sp_, *n*_st_) (Fig. 2a,b). In the homogeneous condensate, *n*_st_ and *n*_sp_ are strongly correlated (Fig. 2a), consistent with a nearly uniform dense phase with local density fluctuations. In contrast, clients in the heterogeneous condensate occupy two distinct microenvironments corresponding to spacer-rich and sticker-rich regions (Fig. 2b). Notably, within the sticker-rich microenvironment, access to the interior of sticker clusters depends strongly on client size: small clients can enter the cores of sticker-rich domains, whereas larger clients are progressively excluded because the insertion cost into these densely packed regions grows fast and nonlinearly with size [22, 23]. As a result, larger clients are increasingly localized near the periphery of sticker clusters (Fig. 2b, inset). This size-dependent shift in client localization provides a qualitative explanation for the unusual size dependence of the attractive contribution Δ*G*_a_ in the heterogeneous condensate.

**FIG. 2.**
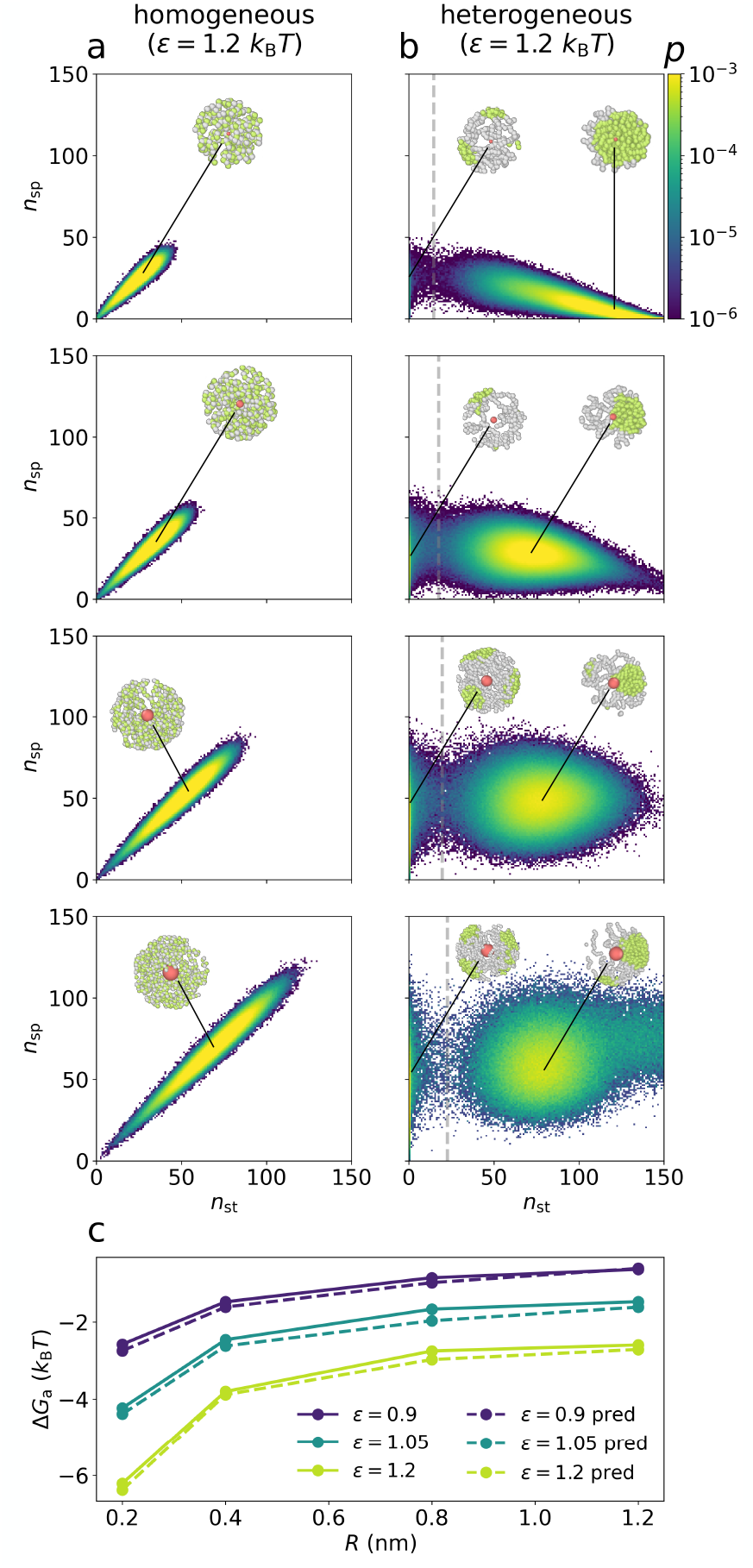
Homogeneous and heterogeneous condensates provide distinct local environments for clients. **a** and **b**, Joint probability distributions of the spacer and sticker coordination numbers, *n*_sp_ and *n*_st_, for clients with radii *R* = 0.2, 0.4, 0.8, and 1.2 nm (top to bottom) at *ϵ*_BC_ = 1.2 *k*_B_*T* in the homogeneous (**a**) and heterogeneous (**b**) condensates. Coordination numbers are calculated within a distance *r*_c_ = *R*+2.5*σ* from the client center. Representative local configurations are shown as insets. For the heterogeneous condensate (**b**), the gray line in each panel separates the spacer-rich (left) and sticker-rich (right) microenvironments and is defined at the local minimum in the probability density along *n*_st_. **c**, Attractive contribution to the transfer free energy, Δ*G*_a_, calculated directly from simulations (solid lines) and predicted from Eq. (2) using the measured occupancies of the two microenvironments (dashed lines). See Methods for details of the simulations and analysis.

To test whether the redistribution quantitatively accounts for the attractive contribution, we further derived a relation between Δ*G*_a_ and the relative occupancies of the two microenvironments. For the parameter regime explored in our simulations, it takes the form

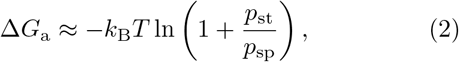

where *p*_st_ and *p*_sp_ are the probabilities of occupying the sticker-rich and spacer-rich states at a given *R* and *ϵ*_BC_, respectively (see Methods for the derivation). As shown in Fig. 2b, the occupancy of the sticker-rich state decreases with increasing client size, which lowers the ratio *p*_st_*/p*_sp_ and thereby |Δ*G*_a_| (Fig. 2c). The change in the attractive contribution, in turn, alters the total transfer free energy and hence client partitioning. Thus, spatial heterogeneity can shape molecular selectivity by modulating the relative occupancies of clients in different dense-phase microenvironments.

### B. Spatial heterogeneity gives rise to heterogeneous transport dynamics

The distinct local environments identified above are also likely to affect the dynamics of molecules within the dense phase. In the heterogeneous condensate, clients can sample both spacer-rich and sticker-rich regions, raising the possibility that their motion alternates between states with different mobilities. To examine this, we performed separate simulations containing only the bulk dense phase of the homogeneous and heterogeneous condensates in periodic cubic boxes, eliminating interfacial effects. The same set of clients with sizes and affinities considered above was then introduced into both dense phases. See Methods for simulation details.

We first tracked the client trajectories and calculated the time- and ensemble-averaged mean-squared displacement (MSD) as a function of lag time Δ*t*. As shown in Fig. 3a, in the homogeneous condensate, clients exhibit essentially normal diffusion over all examined lag times, whereas in the heterogeneous condensate the motion of most clients is subdiffusive at short to intermediate lag times and only approaches normal diffusion at longer lag times, except for the few small and weakly adhesive clients. We further obtained the anomalous diffusion exponent *α* from the logarithmic slope of the MSD, which ranges from 0.7 to 0.9 for most clients in the heterogeneous condensate at short to intermediate lag times (Fig. S2).

**FIG. 3.**
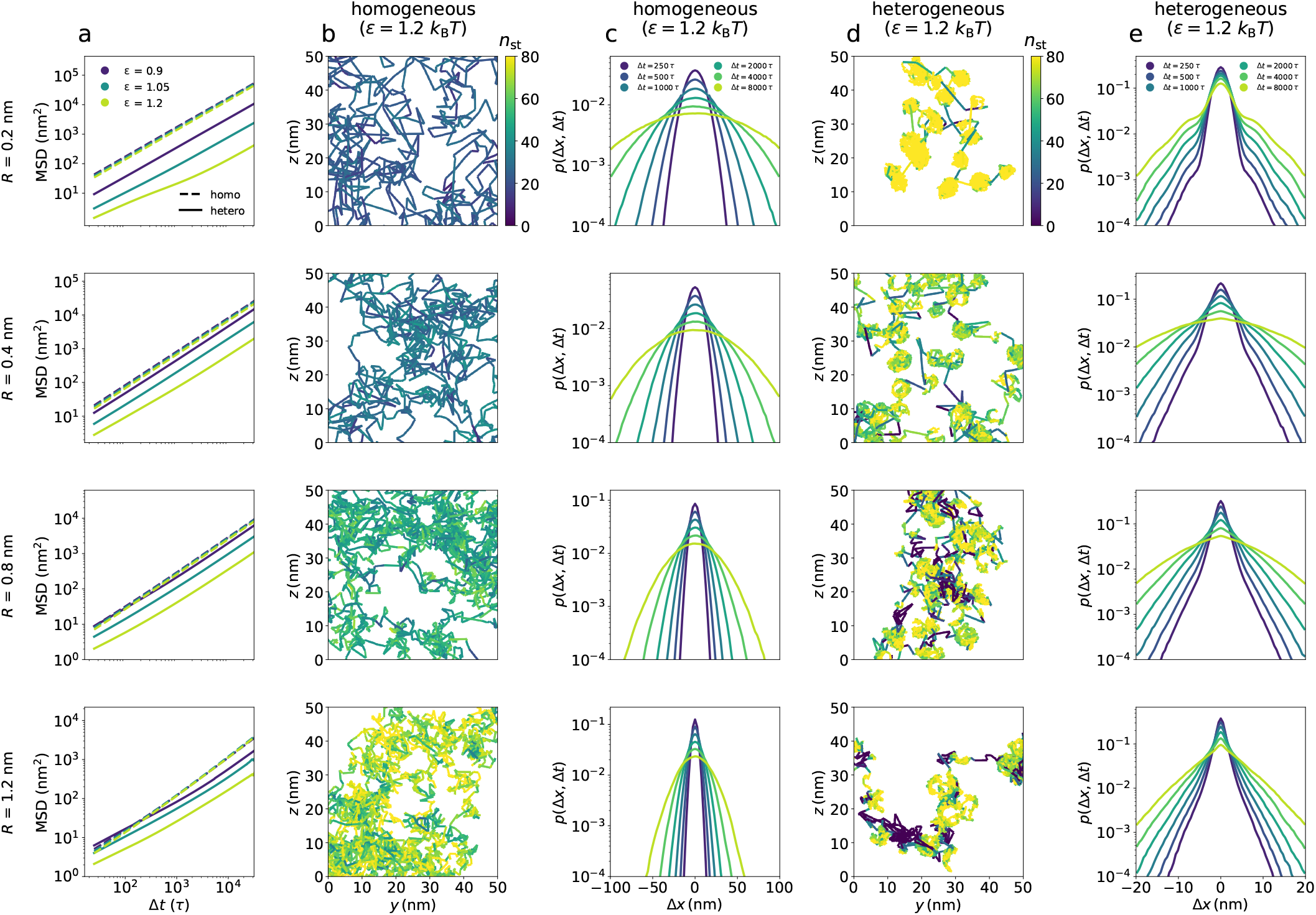
Mesoscale heterogeneity gives rise to heterogeneous client dynamics. Client radii are *R* = 0.2, 0.4, 0.8, and 1.2 nm from top to bottom. **a**, Time and ensemble averaged MSD as a function of lag time Δ*t* at client-sticker affinities *ϵ*_BC_ = 0.9, 1.05, and 1.2 *k*_B_*T* . Δ*t* is given in units of *τ*, where *τ* is the velocity relaxation time in the Langevin dynamics. Dashed and solid lines are for the homogeneous and heterogeneous condensates, respectively. **b** and **d**, Representative client trajectories in the homogeneous (**b**) and heterogeneous (**d**) condensates at *ϵ*_BC_ = 1.2 *k*_B_*T*, colored by the instantaneous sticker coordination number *n*_st_. **c** and **e**, Displacement distributions *p*(Δ*x*, Δ*t*) at lag times Δ*t* = 250, 500, 1000, 2000, 4000, and 8000 *τ* for the homogeneous (**c**) and heterogeneous (**e**) condensates at *ϵ*_BC_ = 1.2 *k*_B_*T* .

The observed subdiffusion suggests that mesoscale organization imposes local constraints on client motion. To visualize these constraints, we plotted representative trajectories for all client sizes at a relatively strong sticker affinity, *ϵ*_BC_ = 1.2 *k*_B_*T*, with each trajectory colored by the instantaneous number of nearby stickers, *n*_st_ (Fig. 3b,d). In the homogeneous condensate, trajectories show fairly uniform *n*_st_ and mobility for a given client size (Fig. 3b). In contrast, trajectories in the heterogeneous condensate consist of locally confined regions connected by more mobile excursions, which are associated with distinct values of *n*_st_ (Fig. 3d). For small clients, localized motion is predominantly with high *n*_st_, corresponding to transient trapping in sticker-rich regions. For larger clients, however, localized motion also occurs at low *n*_st_, corresponding to trapping within cages formed by spacer subchains.

To further characterize client dynamics, we examined the full displacement distributions using the self-part of the van Hove function [38],

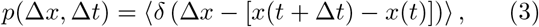

which gives the probability density of a client displacement Δ*x* over a lag time Δ*t*, with *p* averaged over *x, y*, and *z* directions. In the homogeneous condensate, the displacement distributions are essentially Gaussian over all examined lag times (Fig. 3c), as quantified by a near-zero non-Gaussian parameter (Fig. S3). In contrast, the displacement distributions in the heterogeneous condensate are strongly non-Gaussian and vary significantly with client size (Fig. 3e and Fig. S3). For the smallest clients, the distributions exhibit a sharp peak at small displacements and a broad tail at large displacements, which persist even at the longest lag times. As client size increases, the small-displacement peak becomes less pronounced and the distributions develop approximately exponential decay regimes over a range of displacements across multiple lag times.

The non-Gaussian, size-dependent displacement distributions in the heterogeneous condensate reflect the distinct modes of motion observed in the trajectories. For small clients, the sharp peaks at small displacements reflect intermittent trapping in sticker-rich domains, whereas broad tails arise from fast diffusion after escape into spacer-rich domains [39–41]. For larger clients, exclusion from the interiors of sticker-rich domains leads them to sample a broader range of local environments that differ in sticker and spacer densities. The resulting variation in local mobility likely underlies the observed exponential decays in their displacement distributions [40, 42]. Interestingly, even inert clients without sticker attraction exhibit similar exponential regimes (Fig. S4). Overall, these results show that mesoscale organization gives rise to heterogeneous client dynamics by creating distinct local environments with different mobilities.

## III. DISCUSSION

Motivated by recent experimental and computational studies revealing spatially heterogeneous internal organization in biomolecular condensates [26–34], we investigated how such mesoscale organization influences client partitioning and transport. Our simulations show that the homogeneous and heterogeneous condensates exhibit markedly different client partitioning profiles and transport dynamics, because the heterogeneous condensate exposes clients to distinct local microenvironments in a size-dependent manner. These microenvironments are also associated with different mobilities and constraints, giving rise to transient client subdiffusion and non-Gaussian displacement distributions. Our results therefore identify mesoscale organization as an important physical factor controlling both molecular selectivity and transport in biomolecular condensates.

Our simulations provide one specific realization of how mesoscale heterogeneity can shape client partitioning. In our model, sticker-rich domains are also regions of high polymer density. The resulting insertion cost increasingly excludes larger clients from the interiors of sticker-rich domains, causing them to localize near the domain periphery. Although this size dependence may reflect specific features of our minimal model, the overall conclusion is more general: spatial heterogeneity creates local microenvironments that can differ in density, composition, and interaction chemistry, thereby generating distinct local free-energy landscapes for different clients. As a result, client partitioning depends not only on the overall properties of the dense phase, but also on which microenvironments are accessible and energetically favorable to a given client. Different clients can therefore preferentially occupy different regions within the same condensate, or in some cases accumulate at the interfaces between them.

Heterogeneous internal organization could have important consequences for biochemical reactions within condensates. Condensates are commonly thought to regulate reactions by enriching or excluding reactants relative to the surrounding dilute phase [5, 43]. Mesoscale heterogeneity could introduce a further level of compartmentalization, in which enzymes, substrates, regulators, and products preferentially occupy different microenvironments within the same condensate. Such internal organization could enhance reactions by colocalizing reaction partners, suppress them by spatially separating reactants, or organize different steps of a multistep reaction pathway into distinct regions, analogous to the spatially organized RNA processing in the multiphasic nucleolus [44, 45]. Interfaces between microenvironments could also provide additional reaction zones if particular molecules preferentially accumulate there. As with reactions at condensate surfaces [46, 47], such interfacial localization could modify molecular search kinetics [48], while the large total area of internal interfaces could provide more opportunities for interface-mediated chemistry.

Spatial heterogeneity can also influence biochemical activity through its effects on molecular transport. Our simulations show clients sampling local environments with different mobilities and constraints, suggesting that molecular encounters and residence times within heterogeneous condensates cannot necessarily be described by a single effective diffusion coefficient [49]. Consequently, reaction kinetics may depend not only on where reactants are localized, but also on how they move within and between different microenvironments.

Beyond regulating molecular partitioning and transport, mesoscale organization may itself be dynamically coupled to biochemical activity, similar to the coupling of structure and function observed at larger scales in the nucleolus [45, 50]. Local reactions can change molecular binding states, valence, charge, or composition, thereby modifying the interactions that maintain distinct microenvironments within a condensate. These changes could in turn alter client localization and mobility, feeding back on subsequent reactions and molecular transport. This feedback loop could thus allow the internal organization of condensates to reorganize dynamically as biochemical activity changes. Understanding the coupling between mesoscale organization and biochemical activity can therefore be important for connecting condensate structure to its cellular function.

With continuing advances in experimental methods, condensate structure at mesoscopic length scales is becoming increasingly accessible to direct measurement. We hope that this work will motivate further efforts to connect structural heterogeneity to molecular partitioning, transport, and biochemical function in biological condensates.

## IV. METHODS

### A. Coexistence simulations of scaffold–client systems

We performed coarse-grained molecular dynamics simulations using LAMMPS [51, 52]. Each polymer consists of 128 monomers with 64 spacer beads of type A and 64 sticker beads of type B, connected by harmonic bonds

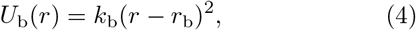

where 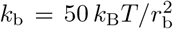 is the spring constant and *r*_b_ = 0.38 nm is the equilibrium bond length, corresponding to the average distance between *α*-carbons of adjacent amino acids in an IDR. Both bead types have diameter *σ* = 0.6 nm.

Sticker-sticker (B-B) pairs interact via an attractive Lennard-Jones (LJ) potential,

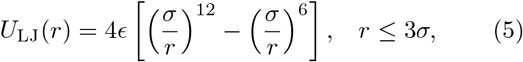

where *ϵ* = *ϵ*_BB_ = 0.84 *k*_B_*T* . The potential was shifted to zero at the cutoff distance. Spacer-spacer (A-A) and spacer-sticker (A-B) pairs repel each other via a WeeksChandler-Andersen (WCA) potential,

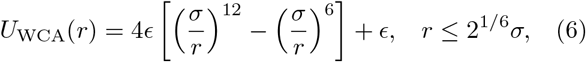

where *ϵ* = *ϵ*_AA_ = *ϵ*_AB_ = 0.84 *k*_B_*T* .

Client particles of type C were modeled as spherical beads of radius *R* = 0.2, 0.4, 0.8, 1.2 nm. Their interactions with polymer beads were described by a radially shifted LJ potential implemented using `lj/expand` in LAMMPS,

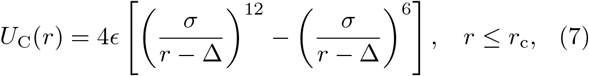

where the shift Δ accounts for the radius of the client particle. For particle-sticker (B-C) interactions, Δ_BC_ = *R* − *σ/*2 and the cutoff distance *r*_c_ is 3*σ* beyond this shift, i.e., *r*_c_ = *R*+ 2.5*σ* from the particle center. The particle-sticker affinity was varied over *ϵ*_BC_ = 0.9, 1.05, 1.2 *k*_B_*T* . For particle-spacer (A-C) and particle-particle (C-C) interactions, Δ_AC_ = *R* − *σ/*2 and Δ_CC_ = 2*R* − *σ*, respectively, and the cutoff is 2^1*/*6^*σ* beyond the shift, so that these interactions are purely repulsive. The repulsive interaction strengths for A-C and C-C were set equal to the corresponding particle-sticker affinity, *ϵ*_AC_ = *ϵ*_CC_ = *ϵ*_BC_. For each interaction type, the potential was shifted to zero at its cutoff distance.

To examine how condensate internal organization affects client partitioning and dynamics, we used two scaffold sequences. In the alternating sequence (*A*_1_*B*_1_)_64_, spacers and stickers alternate along the chain, so that every spacer is adjacent to a sticker. In the blocky sequence (*A*_16_*B*_16_)_4_, the chain is composed of 4 repeating units of 16 spacers followed by 16 stickers.

For each sequence, we simulated 200 polymers and 50 client particles in a 120 nm *×* 20 nm *×* 20 nm box with periodic boundary conditions. To promote the formation of a single dense condensate, polymers were initially confined in the region |*x*| *< x*_wall_ = 23.8 nm by two repulsive LJ walls, corresponding to an initial polymer volume fraction of *ϕ*_init_ = 0.125 in the confined region. The volume fraction was calculated by approximating monomers as spherical beads with a correction to account for overlap between adjacent bonded beads. Client particles were initialized in the dilute region, |*x*| *> x*_wall_, and were allowed to diffuse freely throughout the simulation box at all times.

The time evolution of all beads was simulated using Langevin dynamics [53],

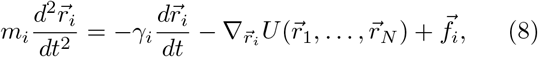

where *m*_*i*_ is the mass of bead *i, γ*_*i*_ is its friction coefficient, 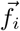 is the random thermal force on bead *i*, and the potential energy *U* is the total potential energy of the system. We used temperature *T* = 300 K, velocity relaxation time *τ* = *m*_*i*_*/γ*_*i*_ = 1 ns, and time step *dt* = 0.01 ns. Bead masses were assigned to reproduce the Stokes-Einstein diffusion coefficient *D*_*i*_ = *k*_B_*T/*(6*πηr*_*i*_) of each bead in water (*η* = 10^−3^ kg/m/s), giving *m*_*i*_ = 6*πηr*_*i*_*τ* .

Equilibration was achieved in three stages. The initial random configurations were first relaxed for 2 *×* 10^6^ steps using soft repulsive interactions between nonbonded beads to eliminate initial steric overlaps. The full pair interactions in Eqs. (5), (6), and (7) were then introduced, and the system was evolved for 1 *×* 10^7^ steps with the confinement walls in place to allow condensate formation. The confinement walls were subsequently removed, and the system was equilibrated for an additional *N ×* 10^7^ steps to allow the dilute phase to form and the condensate to relax further, where *N* = 6 for systems with *ϵ*_BC_ = 0.9, 1.05 *k*_B_*T* and *N* = 11 for systems with *ϵ*_BC_ = 1.2 *k*_B_*T* . Production trajectories were then collected for 5 × 10^7^ steps, with all bead positions saved every 2.5 × 10^5^ steps, yielding 200 frames per replica. 5 replicas for sequence (*A*_1_*B*_1_)_64_ and 25 replicas for sequence (*A*_16_*B*_16_)_4_ were performed to obtain robust statistics.

### B. Determining *ϵ*_BB_ **at matched** *ϕ*_**den**_

To identify the sticker-sticker interaction strength *ϵ*_BB_ that yields the same dense-phase volume fraction for the two sequences, we performed coexistence simulations over a range of *ϵ*_BB_ values at fixed temperature *T* = 300 K. The simulations followed the same protocols as the scaffold-client simulations, except that client particles were not included. The resulting dense- and dilute-phase volume fractions were fit using the law of rectilinear diameters [36]:

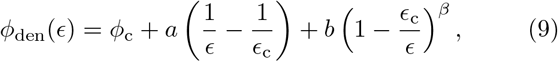

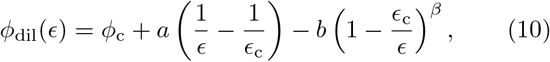

where *ϕ*_c_ is the critical volume fraction, *ϵ*_c_ is the critical interaction strength, *a* sets the diameter slope, *b* sets the amplitude of the coexistence curve splitting, and the exponent *β* = 0.326 corresponds to the 3D Ising universality class. The phase diagrams showed that the two sequences reached the same dense-phase volume fractions, *ϕ*_den_ = 0.19, at *ϵ*_BB_ = 0.84 *k*_B_*T* (Fig. S1). We therefore used this *ϵ*_BB_ value for the comparison between homogeneous and heterogeneous condensates.

### C. Transfer free energy calculation and decomposition

The partition coefficient *P* was computed directly from the simulated particle positions. For each recorded frame, the particle *x*-coordinates were histogrammed using 120 bins of width 1 nm, with the condensate centered in the middle of the box. The histograms were then averaged over all recorded frames and replicas. The dense-phase concentration *c*_den_ was calculated as the mean particle concentration over bins in the dense phase region, |*x*| ≤ 8 nm, and the dilute-phase concentration *c*_dil_ was calculated as the mean particle concentration over bins in the dilute phase region, |*x*| ≥ 30 nm. Both regions lie well away from the condensate interface. The partition coefficient was then obtained as *P* = *c*_den_*/c*_dil_, and the corresponding transfer free energy was computed as Δ*G* = −*k*_B_*T* ln *P*.

The total transfer free energy was decomposed as Δ*G* = Δ*G*_0_ + Δ*G*_a_, where Δ*G*_0_ is the purely repulsive contribution and Δ*G*_a_ is the attractive contribution. The repulsive part was computed by the method of Widom’s test-particle insertion [54, 55]. Specifically, hard-core test particles of radius *R* were inserted at random positions in the dense phase region, |*x*| ≤ 8 nm, and Δ*G*_0_(*R*) = − *k*_B_*T* ln *p*, where *p* is the probability of successful insertion without overlapping with polymer beads. The attractive contribution was then obtained as Δ*G*_a_ = Δ*G* − Δ*G*_0_.

### D. Quantification of microenvironment

We quantified the local environment of each particle by counting the number of spacer beads, *n*_sp_, and sticker beads, *n*_st_, whose centers lie within a cutoff distance *r*_c_ = *R* + 2.5*σ* from the particle center. This cutoff matches the cutoff distance of the particle-sticker interaction in Eq. (7). At each recorded frame, *n*_sp_ and *n*_st_ were computed using `compute coord/atom` in LAMMPS. To restrict the analysis to the dense phase, we only included client particles located in the dense phase region, |*x*| ≤ 8 nm, away from the condensate interface. The instantaneous coordination numbers (*n*_sp_, *n*_st_) were then histogrammed and normalized to obtain the joint probability density *p*(*n*_sp_, *n*_st_), which is shown on a logarithmic scale in Fig. 2. The microenvironments were classified as spacer-rich or sticker-rich using a decision boundary defined by a constant value of *n*_st_. The boundary *n*_st_ was chosen at the local minimum of the probability density between the two metastable states. For *ϵ*_BC_ = 1.2 *k*_B_*T* and *R* = 0.2, 0.4, 0.8, 1.2 nm, the corresponding boundaries are *n*_st_ = 14, 17, 19, 22, respectively, which are shown as gray lines in Fig. 2b.

### E. Derivation of Eq. (2)

The transfer free energy for moving a client from the dilute phase to the dense phase depends on its radius *R* and sticker affinity *ϵ* and can be written as

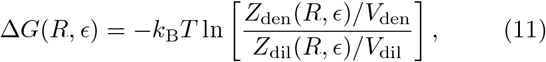

where *Z*_*α*_(*R, ϵ*) is the configurational partition function of the client in phase *α* = den, dil,

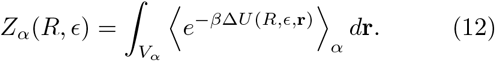

Here, Δ*U*(*R, ϵ*, **r**) is the interaction energy associated with inserting a client at position **r**, ⟨· · · ⟩_*α*_ denotes an ensemble average over scaffold configurations in phase *α*, and *V*_*α*_ is the corresponding phase volume. The attractive contribution to the transfer free energy, Δ*G*_a_(*R, ϵ*) = Δ*G*(*R, ϵ*) − Δ*G*(*R*, 0), is then

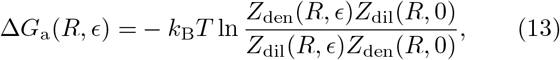

where the phase-volume factors cancel out.

For the heterogeneous condensate, we decompose the dense-phase partition function into contributions from the spacer-rich and sticker-rich microenvironments,

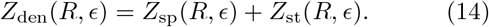

For the parameter regime considered here, we make three approximations. First, *Z*_dil_(*R, ϵ*) ≈ *Z*_dil_(*R*, 0) because client-sticker interactions contribute little in the dilute phase. Second, *Z*_st_(*R*, 0) ≪ *Z*_sp_(*R*, 0) because inert clients are strongly excluded from the sticker-rich microenvironment. Third, *Z*_sp_(*R, ϵ*) ≈ *Z*_sp_(*R*, 0) because clients in the spacer-rich microenvironment make relatively few sticker contacts. Under these approximations, Eq. (13) becomes

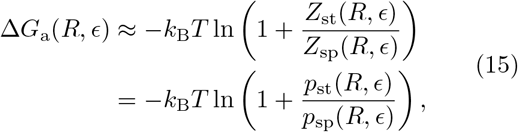

where *p*_*i*_ = *Z*_*i*_*/*(*Z*_sp_ + *Z*_st_) is the probability that a client occupies microenvironment *i*. Thus, within these approximations, the attractive contribution to the transfer free energy is determined by the relative occupancies of the sticker-rich and spacer-rich microenvironments.

### F. Characterization of client dynamics

To characterize client dynamics within the homogeneous and heterogeneous condensates while avoiding interfacial boundary effects, we performed separate simulations containing only the bulk dense phase in a fully periodic cubic box of side length *L* = 23.8 nm. Each simulation contained 216 polymers initialized on a 6 *×* 6 *×* 6 grid, resulting in a target dense-phase volume fraction *ϕ*_den_ = 0.19. 10 client particles were added for *in silico* single-molecule tracking. The interaction potentials and Langevin thermostat parameters were the same as those used in the coexistence simulations. The system was equilibrated using the procedure described in the first Methods subsection with *N* = 6 for all cases. During production, particle positions and local environments (*n*_sp_ and *n*_st_) were saved every 2500 steps (25 *τ*), yielding a total of 2 *×* 10^4^ recordings. 5 and 15 independent replicas were performed for the homogeneous and heterogeneous condensates, respectively.

The time and ensemble averaged MSD was calculated

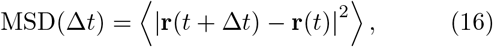

where the time average was computed using an FFT-based algorithm [56] for each client trajectory, with lag times up to one quarter of the trajectory length. The ensemble average was then taken over all client trajectories.

The effective anomalous diffusion exponent *α* was obtained from the local logarithmic slope of the MSD,

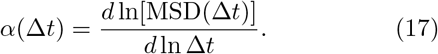

The displacement distributions were constructed from the single-coordinate displacements Δ*x* = *x*(*t* + Δ*t*) − *x*(*t*), with statistics pooled over the *x, y*, and *z* directions as well as all available time origins and client particles. The resulting distributions were normalized to obtain the self-part of the van Hove function *p*(Δ*x*, Δ*t*). The nonGaussian parameter shown in Fig. S3 was calculated as

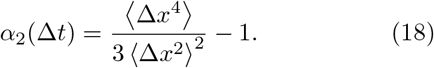

## ACKNOWLEDGEMENTS

We thank Ned S. Wingreen for valuable comments and suggestions on the manuscript. V.G. and Y.Z. were supported by a startup fund at Johns Hopkins University, NIH Award R35GM162296, and the Alfred P. Sloan Foundation through a Sloan Research Fellowship to Y.Z. (FG-2025-25076). This work was carried out at the Advanced Research Computing at Hopkins (ARCH) core facility (rockfish.jhu.edu), which is supported by the National Science Foundation (NSF) grant number OAC1920103.

