## Supplemental Material for "Mesoscale Heterogeneity Shapes Molecular Selectivity and Transport in Biomolecular Condensates"

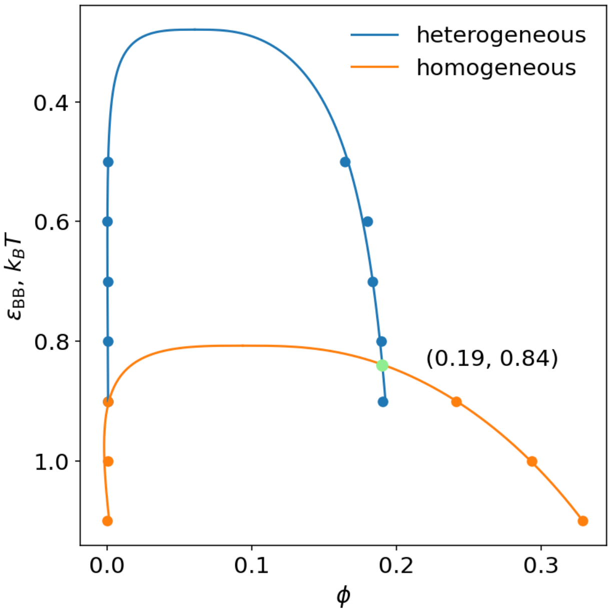


FIG. S1: Phase diagrams for the heterogeneous (blue) and homogeneous (orange) condensates. Points show the dilute-phase ($|x| > 30$ nm) and dense-phase ($|x| < 8$ nm) volume fractions $\phi$ measured at each interaction strength $\varepsilon_{\mathrm{BB}}$ in simulations. Lines are fits using the law of rectilinear diameters. The green point marks the intersection of the two phase boundaries at $(\phi,\, \varepsilon_{\mathrm{BB}}) = (0.19,\ 0.84\, k_{\mathrm{B}}T)$.

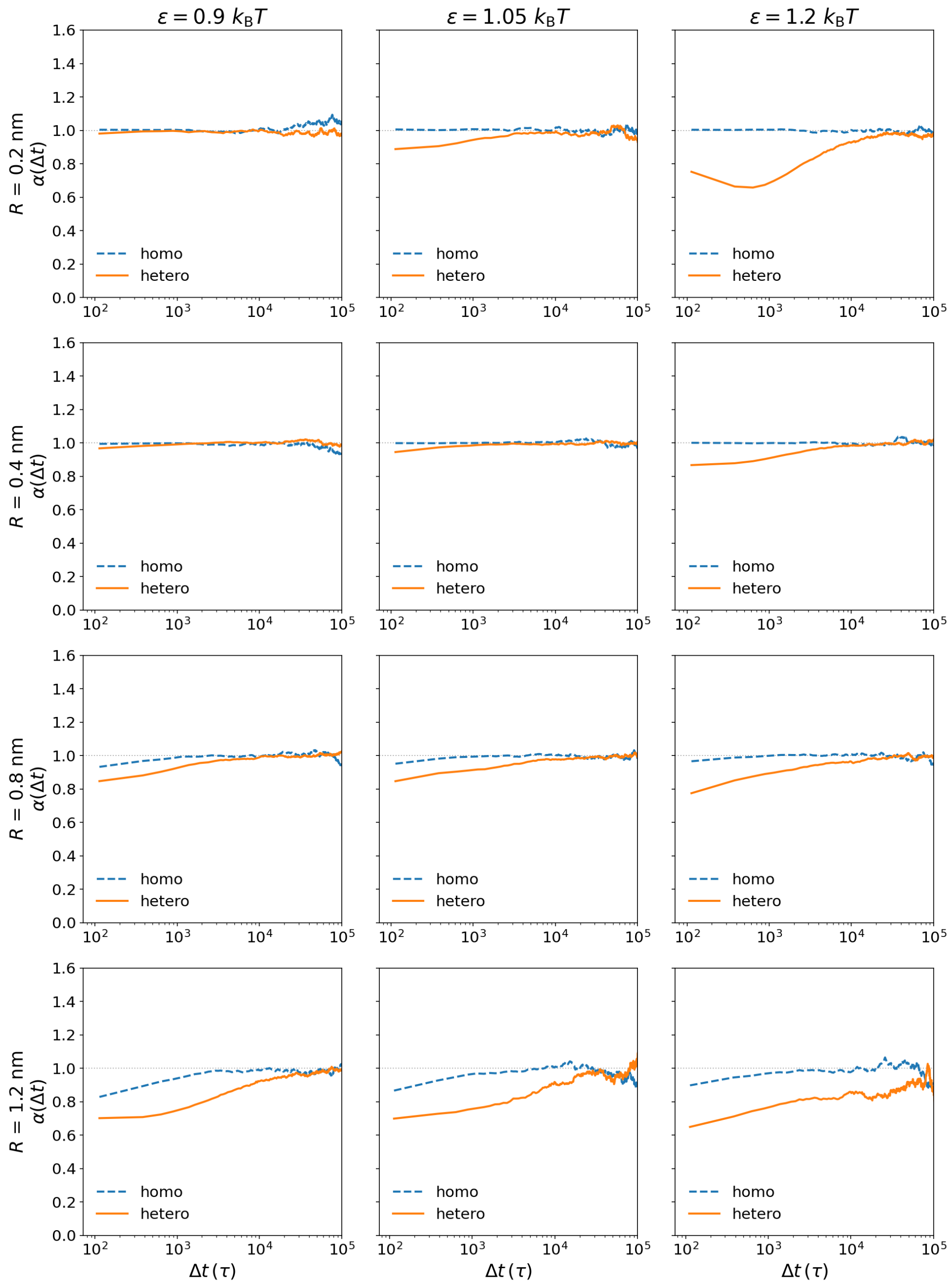


FIG. S2: The anomalous exponent $\alpha(\Delta t) \equiv d\ln[\mathrm{MSD}(\Delta t)]/d\ln(\Delta t)$ as a function of the lag time $\Delta t$ for client motion in the homogeneous (blue) and heterogeneous (orange) condensates. Results are shown for client radii $R \in \{0.2, 0.4, 0.8, 1.2\}$ nm and interaction strengths $\varepsilon_{\mathrm{BC}} \in \{0.9, 1.05, 1.2\}\ k_\mathrm{B}T$. To reduce fluctuations arising from numerical differentiation, $\alpha(\Delta t)$ was obtained by fitting $\ln[\mathrm{MSD}(\Delta t)]$ versus $\ln(\Delta t)$ in non-overlapping windows of 10 data points. The dotted horizontal line indicates the expected value $\alpha = 1$ for normal diffusion.

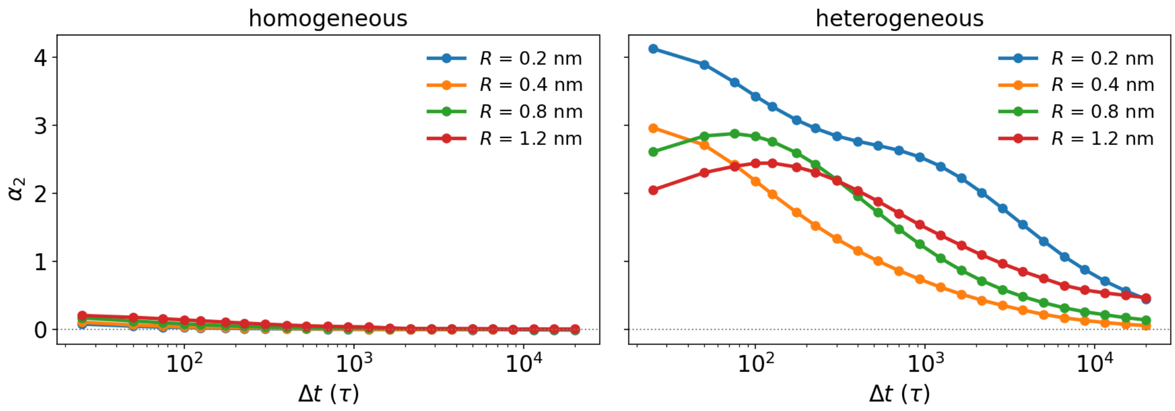


FIG. S3: The non-Gaussian parameter $\alpha_2(\Delta t) \equiv \langle \Delta x^4 \rangle / (3\langle \Delta x^2 \rangle^2) - 1$ for the displacement distribution of client motion in the homogeneous (left) and heterogeneous (right) condensates. Results are shown for client radii $R \in \{0.2, 0.4, 0.8, 1.2\}$ nm and an interaction strength $\varepsilon_{\mathrm{BC}} = 1.2\, k_{\mathrm{B}}T$.

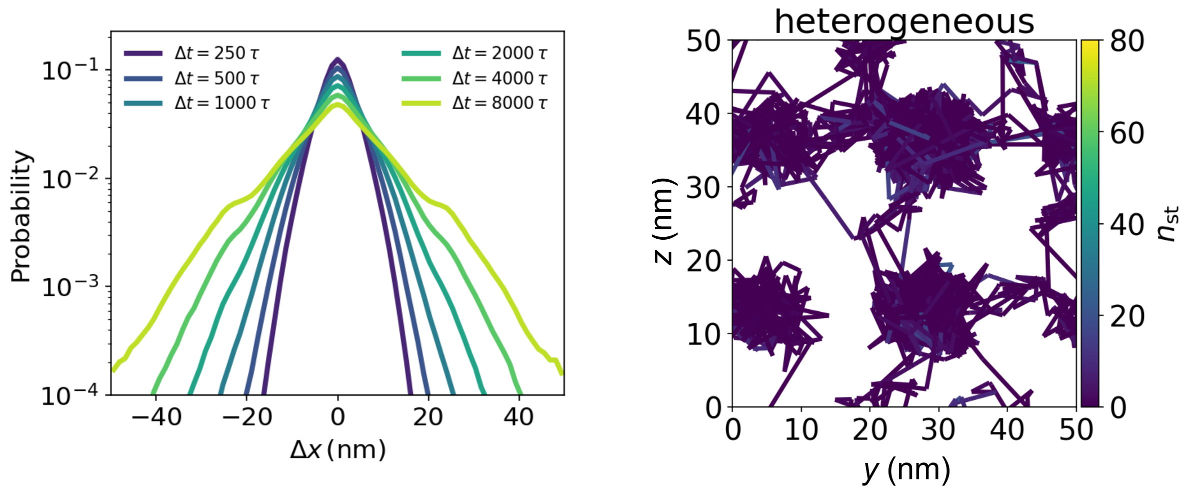


FIG. S4: Displacement distributions (left) and a representative trajectory (right) of an inert probe particle ($R = 1.2$ nm, $\varepsilon_{\mathrm{BC}} = 0\, k_{\mathrm{B}}T$) diffusing in the bulk of a heterogeneous condensate.
